# Spatial Autocorrelation Aware Resampling Improves Cell-Cell Interaction Inference in Spatial Transcriptomics Data

**DOI:** 10.64898/2026.07.06.736800

**Authors:** Parth Khatri, Michael A Newton, Christina Kendziorski, Huy Q. Dinh

**Affiliations:** McArdle Laboratory for Cancer Research, Department of Oncology, University of Wisconsin-Madison; Department of Biostatistics and Medical Informatics, University of Wisconsin-Madison; Department of Statistics, University of Wisconsin-Madison

## Abstract

Spatial transcriptomics has enabled finer-grained analyses of cell-cell interactions through the co-expression of ligands and their cognate receptors, thereby accounting for the spatial constraints of signaling. However, existing methods employ random permutations or analytic calculations to assess statistical significance, neither of which accounts for spatial autocorrelation, a common property of spatially resolved data. Here, we introduce SOAAR (Spatial Omics Autocorrelation-Aware Resampling), a statistical method for testing gene-gene correlations in spatial data that maintains spatial gene-level autocorrelation in resampled datasets used to generate null distributions. SOAAR uses spatial map patterns to decompose autocorrelation. The associations between gene expression and autocorrelation patterns are then randomized to construct resampled datasets used for evaluating significance testing. We showed that SOAAR maintains gene-level spatial autocorrelation and yields a lower false-positive rate than random permutations across varying degrees of gene-level spatial autocorrelation in simulation studies. In a 10X Visium dataset from 10 HNSCC patients treated with immunotherapy, SOAAR filters out low-confidence interactions that were present in only individual samples or in fewer than 3 samples. That led to the identification of a consistent signature of T-cell recruitment in Responder patients and a resistance signature driven by angiogenesis and tumor cell proliferation in Non-Responders. Similar trends were observed in a larger cohort of 23 patients profiled with single-cell spatial CosMX SMI data, revealing immune cell interactions in response to immunotherapy. Overall, SOAAR provides a more calibrated framework for testing spatial correlation, grounded in spatial statistics. Future developments will seek to link localized correlation patterns to downstream changes in biological pathways, thereby informing biomarker and therapeutic target discovery.

## BACKGROUND

Cell-cell communication is crucial for tissue and organism-level functioning. During disease, these communication networks can become dysregulated or rewired, providing potential biomarkers for diagnosis or therapeutic targets [add a citation]. The expansion of single-cell transcriptomics technologies has enabled the systematic study of cell-cell communication via thousands of ligand-receptor interactions between specific cell populations. Methods like CellPhoneDB [1], CellChat [2], and NATMI [3] enable cell-cell communication inference by identifying pairs of cell populations where one has upregulated ligand expression, and another has upregulated expression of the ligand’s cognate receptor [4]. The removal of cells from their tissue context, as in scRNA-Seq data, reduces the efficacy of these inferences due to the loss of cell-cell proximity. The development of spatial transcriptomics (ST) can address that limitation through profiling of gene expression with spatial context. At the same time, computational methods for scRNA-seq data do not accommodate spatial data. Notably, most scRNA-seq methods require prior clustering, based on gene expression similarity, before the cell-cell communication inference step [5]. In spatial omics data, clustering just based on gene expression loses spatial information, preventing the identification of spatially localized interactions. Thus, frameworks for studying cell-cell communication in spatial transcriptomics data require a different approach that does not rely on prior cell-type clustering.

Inferring cell-cell communication in spatial transcriptomics data is an open question, and many methods have been developed to address it. Graph learning [6–9], optimal transport [10], dimensionality reduction [11], and spatial correlation-based [12, 13] approaches have all been applied to address this question. The spatial correlation-based approaches typically calculate bivariate Moran’s I [14], a measure of proximal spatial co-localization between two features, and then generate a null distribution to assess statistical significance. SpatialDM [12] permutes location identifiers (barcodes in sequencing-based ST or cell IDs in fluorescence-based ST) to determine statistical significance, while MERINGUE [13] permutes cell type labels to identify enriched ligand-receptor interactions between cell types in single-cell resolution ST. While these approaches may be appropriate for testing whether two genes have independent spatial distributions, they are insufficient to test whether two genes with individual spatial structures are co-localized. Motivated by the field of spatial statistics, univariate Moran’s I [15] has been used to characterize the spatial structure of a feature, quantified by its spatial autocorrelation. Existing methods to test the significance of bivariate Moran’s I [12, 13] assume that features are not spatially autocorrelated. We hypothesize that this assumption leads to an underestimation of the variance under the null and an increased number of bivariate correlations deemed statistically significant. To address this gap in existing methods for bivariate spatial correlation testing, we propose Spatial Omics Autocorrelation Aware Resampling (SOAAR) – a hypothesis testing framework for bivariate spatial correlations by preserving each gene’s spatial autocorrelation structure. This enables the construction of surrogate datasets for testing bivariate spatial correlation while addressing potential confounding. SOAAR outperforms existing methods and achieves a substantial reduction in false positives while being adaptive to gene-level spatial autocorrelation, in rigorous simulation settings, and in an application to a dataset of head and neck squamous cell carcinoma (HNSCC) 10X Visium samples from patients treated with immune checkpoint blockade [16].

## RESULTS

### Overview of the SOAAR Framework for bivariate correlation analysis in ST data

SOAAR (**Fig. 1A**) applies Moran Spectral Randomization [17] from spatial ecology to the spatial-weights graph of spatial locations (cells or spots) in ST data as input. In practice, we used Ligand-Receptor pairs to illustrate the method’s performance and application. Maintaining feature-level spatial autocorrelation patterns enables better calibrated testing of spatial correlations, reducing false-positive rates and spurious correlation findings observed in existing methods, thereby facilitating confident identification of spatial interactions.

**Figure 1.**
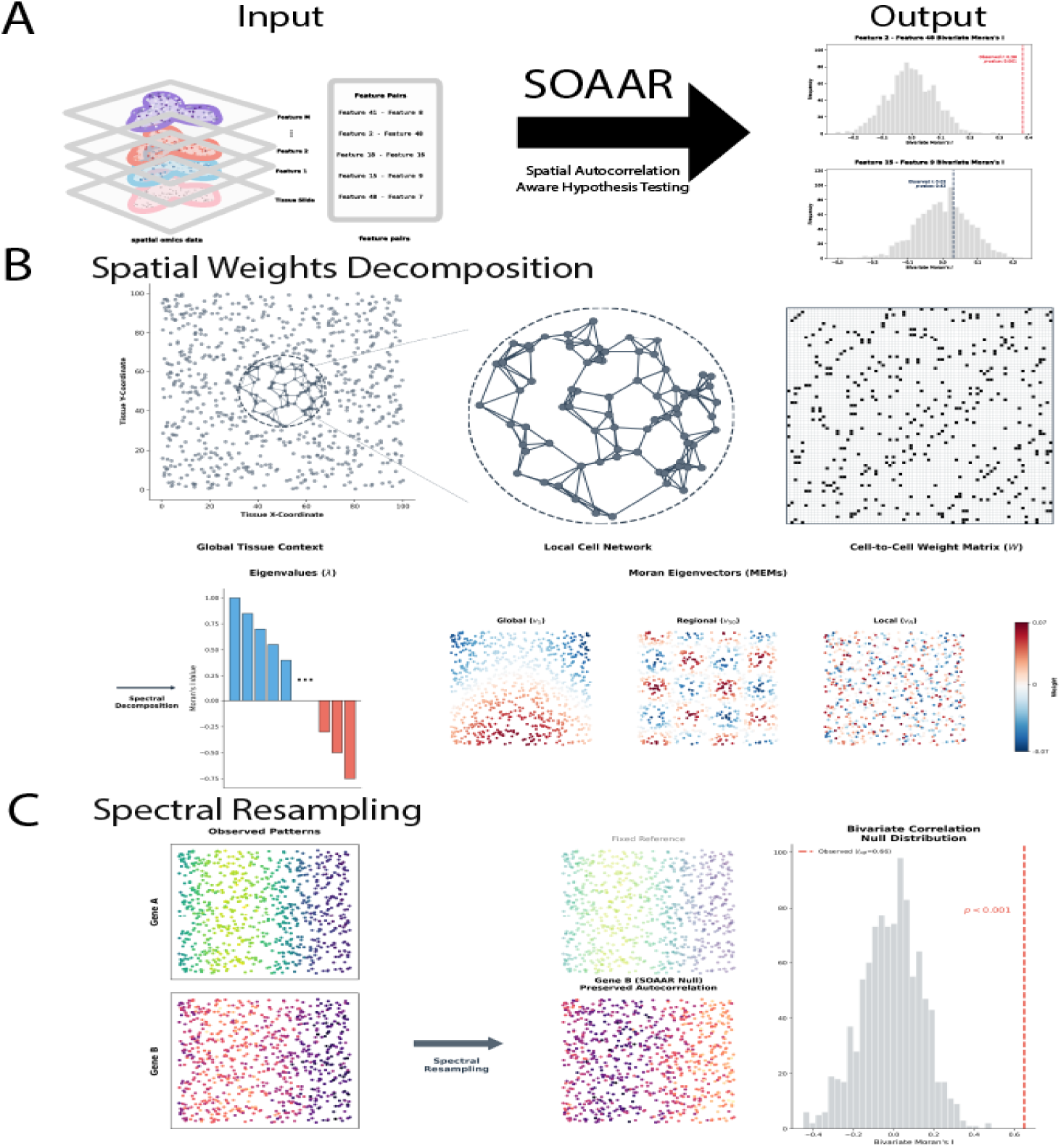
Overview of the SOAAR framework for autocorrelation-aware spatial co-expression testing. A. SOAAR takes as input an AnnData object that contains a spatial expression matrix and coordinates, and a database of interacting features, returning a bivariate Moran’s I statistic for each interacting feature pair and the associated p-values and adjusted p-values. Example histograms illustrate a non-significant pair (top) and a significant pair (bottom). B. SOAAR uses the spatial connectivity between locations to construct a spatial weights matrix, which is then doubly centered and spectrally decomposed into the Moran Eigenvectors (MEMs), a set of spatial autocorrelation patterns spanning global, regional, and local scales (ordered by decreasing eigenvalue). C. SOAAR’s spectral resampling procedure. For a given gene pair (Gene A, fixed reference; Gene B, to be resampled), SOAAR projects Gene B onto the MEM basis and randomizes the spectral coefficients while preserving the power spectral density, generating surrogate expression patterns that maintain the original spatial autocorrelation structure. The resulting set of surrogate bivariate Moran’s I values forms the null distribution to test the observed statistic.

SOAAR’s application of Moran Spectral Randomization starts with the spectral decomposition of the doubly centered user-defined spatial weights matrix (**Fig. 1B**). By default, SOAAR separates interactions into short- and long-range interactions using two different weight matrices: a k-nearest neighbors’ graph and a radial basis function with bandwidth set to twice the smallest pairwise distance. The resulting eigenvectors are the Moran Eigenvectors, spatial patterns that can be used to reconstruct gene expression patterns in the assayed tissues. The associated eigenvalues describe Moran’s I for each of these patterns. From there, interacting genes are then correlated with Moran Eigenvectors, and the spectral randomization is performed using one of three procedures – a singleton procedure, a random pair procedure, or a sequential pair procedure (Methods). SOAAR then outputs a Pandas [18] dataframe that indicates the observed bivariate Moran’s I for each pair as well as an associated p-value, adjusted p-value after the Benjamini-Hochberg correction, and a Boolean significance variable based on the BH p-value with a default cutoff of 0.05. This is stored in the input AnnData [19] object by default, but can be optionally exported as a .csv file. SOAAR’s procedure enables statistical testing with a more comprehensive, spatially aware hypothesis-testing framework that accounts for the spatial structure of univariate gene expression, rather than existing methods that test bivariate spatial correlation under the assumption of random spatial structure at the single-gene level.

SOAAR is implemented in Python and is designed for easy interoperability with scverse software [20] such as AnnData [19], scanpy [21], and squidpy [22]. Sparse matrix operations and the Nystrom approximation enable fast, scalable computations as the number of cells increases, despite computationally intensive steps such as spectral decomposition in the workflow.

### SOAAR preserves spatial autocorrelation, reduces type I error while maintaining statistical power in simulated data

Given that the primary difference between SOAAR’s approach and existing methods lies in the preservation of autocorrelation, we compare the performance of SOAAR and random permutation across different simulation settings to evaluate how SOAAR’s MSR-based randomization approach affects autocorrelation preservation, false-positive rate, and statistical power.

To evaluate how well SOAAR’s different randomization procedures preserve spatial autocorrelation, we simulated normally distributed genes across a range of autocorrelation coefficients, with 500 genes generated per coefficient (Methods). We then evaluated the root mean squared error (RMSE) of the autocorrelation for 1000 surrogates generated by SOAAR’s singleton randomization procedure, random eigenvector pairing procedure, and sequential eigenvector pairing procedure, and a random permutation of values for each simulated gene, like those used in SpatialDM [12] or MERINGUE [13]. Across autocorrelation values, the uniform random permutation showed that it could not preserve spatial autocorrelation (**Fig. 2A**). Even in cases where the coefficient was near 0, the uniform random permutation had a mean RMSE of 0.050. Across all coefficients, the mean RMSE of the random permutation was 0.477 with a variance of 0.0019. When SOAAR’s pairing of eigenvectors was random, the mean RMSE was 0.252 with a variance of 0.0021, showing improvement over the uniform random permutation. Further, in both the SOAAR’s singleton and the sequential eigenvector pairing procedures, the mean RMSE of spatial autocorrelation was 0.0102 and 0.0107, with variances of 0.000015 and 0.000014, respectively. This shows that strictly maintaining the power spectral density using the singleton procedure or approximately maintaining it using the sequential pair procedure both led to very strong preservation of spatial autocorrelation at the single-gene level.

**Figure 2.**
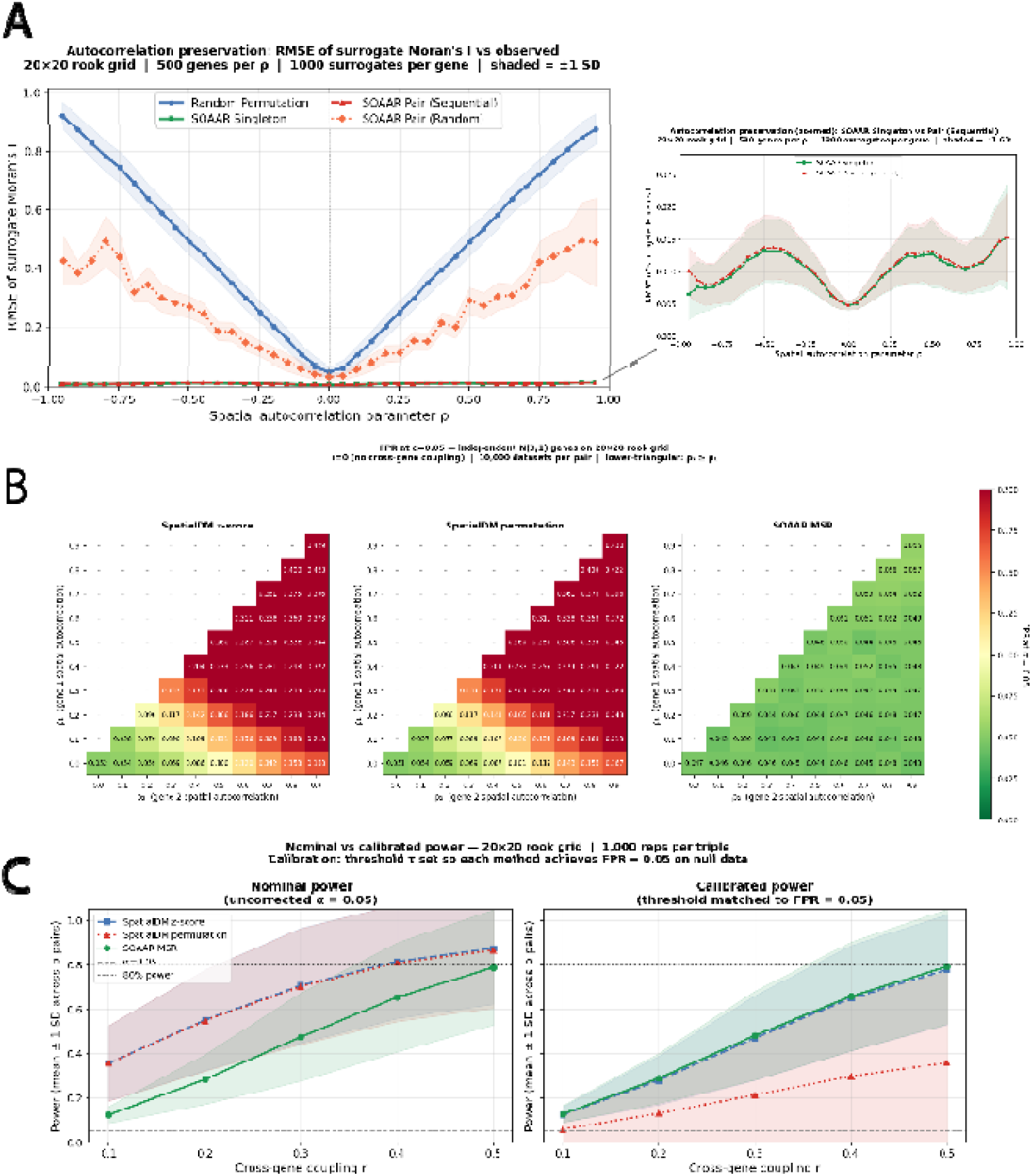
SOAAR preserves spatial autocorrelation, controls false-positive rates, and achieves superior calibrated power compared to SpatialDM on simulated data. A. Root mean squared error (RMSE) between the surrogate and observed Moran’s I across a range of spatial autocorrelation parameters (ρ, −0.95 to 0.95), evaluated on a 20×20 rook grid with 500 simulated genes per ρ and 1,000 surrogates per gene (shading = ±1 SD). To capture the scale of RMSE for SOAAR Singleton and SOAAR Pair (Sequential), a zoomed-in plot is provided. B. False positive rate (FPR) at α = 0.05 under the null (no cross-gene coupling, r = 0) for 10,000 simulated gene pairs per ρ₁,ρ₂ pair on a 20×20 rook grid. C. Nominal (left) and calibrated (right) statistical power as a function of cross-gene coupling strength (r = 0.1–0.5), averaged across autocorrelation regimes (1,000 replicates per ρ₁, ρ₂, r triplet).

After confirming that SOAAR can effectively preserve spatial autocorrelation in its surrogates, we examined its ability to control false-positive rates. Here, we generated simulated gene pairs with differing degrees of positive spatial autocorrelation and no cross-correlation. Negative autocorrelation regimes were excluded in this simulation as negative autocorrelation is rare in spatial data and when it is observed, it tends to have low magnitude close to 0. To compare with SOAAR, we then used SpatialDM [12], a widely used method that employs bivariate Moran’s I to test for statistically significant ligand-receptor interactions in spatial transcriptomics data. For each pair of autocorrelation coefficients, we simulated 10,000 gene pairs and generated 1000 surrogates per gene pair using SpatialDM’s permutation, analytic z-score, and SOAAR’s eigenvector pairing procedure. SpatialDM was able to maintain a false positive rate of 0.05 when ρ_1_=0 and ρ_2_=0 or 0.1 (**Fig. 2B**). However, once the autocorrelation coefficient increases, the false positive rate exceeds 0.05. If both autocorrelation coefficients are greater than or equal to 0.4, the false positive rate was at least 0.2 and increased further as either autocorrelation coefficient increased. In contrast, SOAAR showed consistent control of false positives, with a rate below 0.05 if at least one autocorrelation coefficient was below 0.8. With both autocorrelation coefficients greater than or equal to 0.8, SOAAR’s false positive rate increases to 0.06, which is comparable to the false positive rate of both of SpatialDM’s methods at {(0.1,0.1), (0,0.2)}. These results showed SOAAR’s spatially aware randomization outperforms SpatialDM in controlling Type I error. In contrast, assuming spatially random gene distributions increases the number of false positives, even at modest levels of spatial autocorrelation. This is confirmed by examining the distributions of the p-values: SpatialDM shows p-values clustered at 0 and 1, whereas SOAAR shows a relatively flat distribution resembling a uniform(0,1) distribution (**Fig. S1**), even at extreme degrees of spatial autocorrelation.

Our final simulation tests the statistical power of SOAAR’s sequential eigenvector pairing procedure and SpatialDM’s permutation-based and analytic z-score methods. To do so, we simulated 1,000 gene pairs for each of 15 different autocorrelation pairs with 5 different values of cross-correlation, making 75 total triplets and 75,000 total gene pairs. Across triplets, the nominal power of SpatialDM is higher than SOAAR, especially at lower effect sizes (**Fig. 2C**, left). After the calibration, SOAAR still performs slightly better than SpatialDM’s analytic calculation and substantially better than SpatialDM’s permutation (**Fig. 2C**, right).

Altogether, the results of our simulation studies show SOAAR’s capability to preserve spatial autocorrelation with low error across a spectrum of possible levels of spatial autocorrelation, and to control false positive rates under the null despite additional noise introduced by spatially autocorrelated variables.

### SOAAR identifies immunotherapy-associated cell-cell interactions in 10X Visium data from head and neck squamous cell carcinoma

Next, we evaluate the results of SpatialDM and SOAAR on a dataset of 10 HNSCC tumor samples (see Methods) collected prior to immunotherapy. For each sample, we defined the weight matrix for secreted signaling interactions using a radial basis function kernel with bandwidth equal to twice the minimum spot-to-spot distance of adjacent spots, and the cell-ECM and cell-cell contact interactions using the radial nearest neighbors graph, where r is the maximum spot-to-spot distance of adjacent spots.

Our results show that SOAAR reports substantially fewer interactions in almost every sample (**Fig. 3A**). When combining across samples and considering interactions that are unique to SpatialDM, unique to SOAAR, or shared by both, SOAAR filters out 35.8% of interactions. The exception to this lies in sample ck17-1592, where SOAAR identifies a near equal number of unique interactions to SpatialDM, but the structure of the tissue may be the cause of this, since it is composed of two separate connected components, which may cause some mixing of distinct gene expression from each component in the null surrogates. However, in terms of distribution, the detection rate of SOAAR between responsive and non-responsive samples does not differ (Mann-Whitney p-value = 0.69). Additionally, due to the higher degree of calibration, SOAAR filters out interactions that are identified in three or fewer samples. Most interactions identified by SpatialDM alone are restricted to one or two samples, whereas those SOAAR shares with SpatialDM span multiple interactions (**Fig. 3B**). In terms of statistical power, this reduces the number of potential outliers, thereby prioritizing further biological validation in multi-sample analyses.

**Figure 3.**
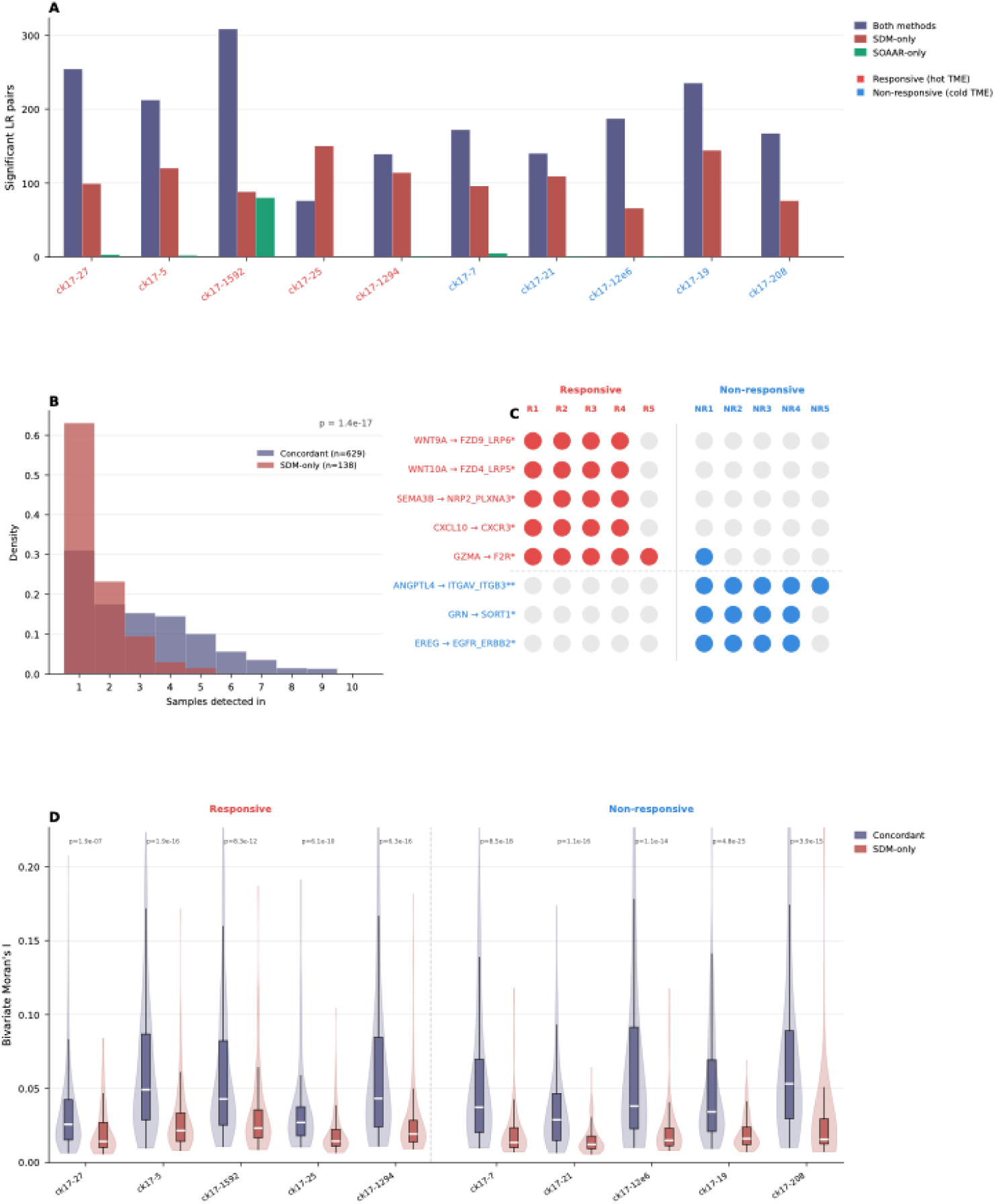
SOAAR reduces spurious ligand-receptor interactions and identifies biologically coherent spatial co-expression patterns associated with pembrolizumab response in HNSCC Visium data. A. Number of significant ligand-receptor pairs detected per sample across 10 Visium HNSCC samples (5 immunotherapy-responsive, 5 non-responsive), broken down by significant interactions called by both methods (concordant), by SpatialDM only (SDM-only), or by SOAAR only. B. Distribution of the number of samples in which each interaction was detected, stratified by concordant and SDM-only interactions (Mann-Whitney p = 1.4×10⁻¹□). C. Heatmap of selected ligand-receptor interactions significantly detected across responsive (R1–R5) and non-responsive (NR1–NR5) samples (* p < 0.05; ** p < 0.01). D. Distribution of bivariate Moran’s I value for concordant interactions versus SDM-only interactions across all 10 samples (per-sample Mann-Whitney p-values shown).

Among the statistically significant Ligand-Receptor pairs identified, we found CXCL10-CXCR3 and GZMA-F2R mostly in Responders (4 out of 5) (**Fig. 3C, Table S1**). CXCL10-CXCR3 is potential evidence of T cell recruitment into the tumor microenvironment, which was identified as a part of the immunotherapy response signature in our and others’ previous work [16]. The GZMA-F2R interaction has been identified as both an interaction between T cells and endothelial cells as a regulator of angiogenesis, and between T cells and tumor cells as a tumor growth suppression mechanism [23, 24]. Among the Non-Responders, ANGPTL4-ITGAV_ITGB3 interactions are commonly associated with angiogenesis, a pro-tumoral feature in the tumor microenvironment [25], GRN-SORT1 associated with expansion of cancer stem cells [26], and EREG-EGFR_HER2, a marker of tumor progression [27, 28]. In terms of effect size, the distribution of bivariate Moran’s I from the shared interactions to Moran’s I called significant by only SpatialDM, the median bivariate Moran’s I shows a substantial difference (**Fig. 3D**). This is in line with our simulation study that SOAAR is a more sensitive test, calling fewer interactions as significant and only from interactions that have a higher Moran’s I.

### SOAAR dissects immune interactions associated with immunotherapy-responsive and non-responsive HNSCC tumor microenvironments in Bruker CosMx SMI data

To explore SOAAR’s applicability to single-cell spatial transcriptomics data, we used a Bruker CosMx SMI dataset comprising 68 samples, represented as fields of view (FOVs) from a tumor microarray made from 23 head and neck squamous cell carcinoma patients treated with immune checkpoint blockade immunotherapy. The CosMx SMI panel includes 986 genes, including 435 ligands and receptors, and 363 immune-related pairs, enabling us to examine interactions between immune cells and other cell types.

The initial number of detected interactions followed a trend similar to that observed in the 10X Visium data: FOVs generally showed that SpatialDM detects more significant interactions unique to the method than SOAAR, while most patients showed overlap across both methods (**Fig. 4A**). Unlike the Visium data, more patients seemed to deviate from that trend, though that could be a function of a larger sample size. Similar to the results from Visium, the interactions identified as significant by SpatialDM alone were also observed in fewer patients, on average, than those identified as statistically significant by both methods (**Fig. 4B**).

**Figure 4.**
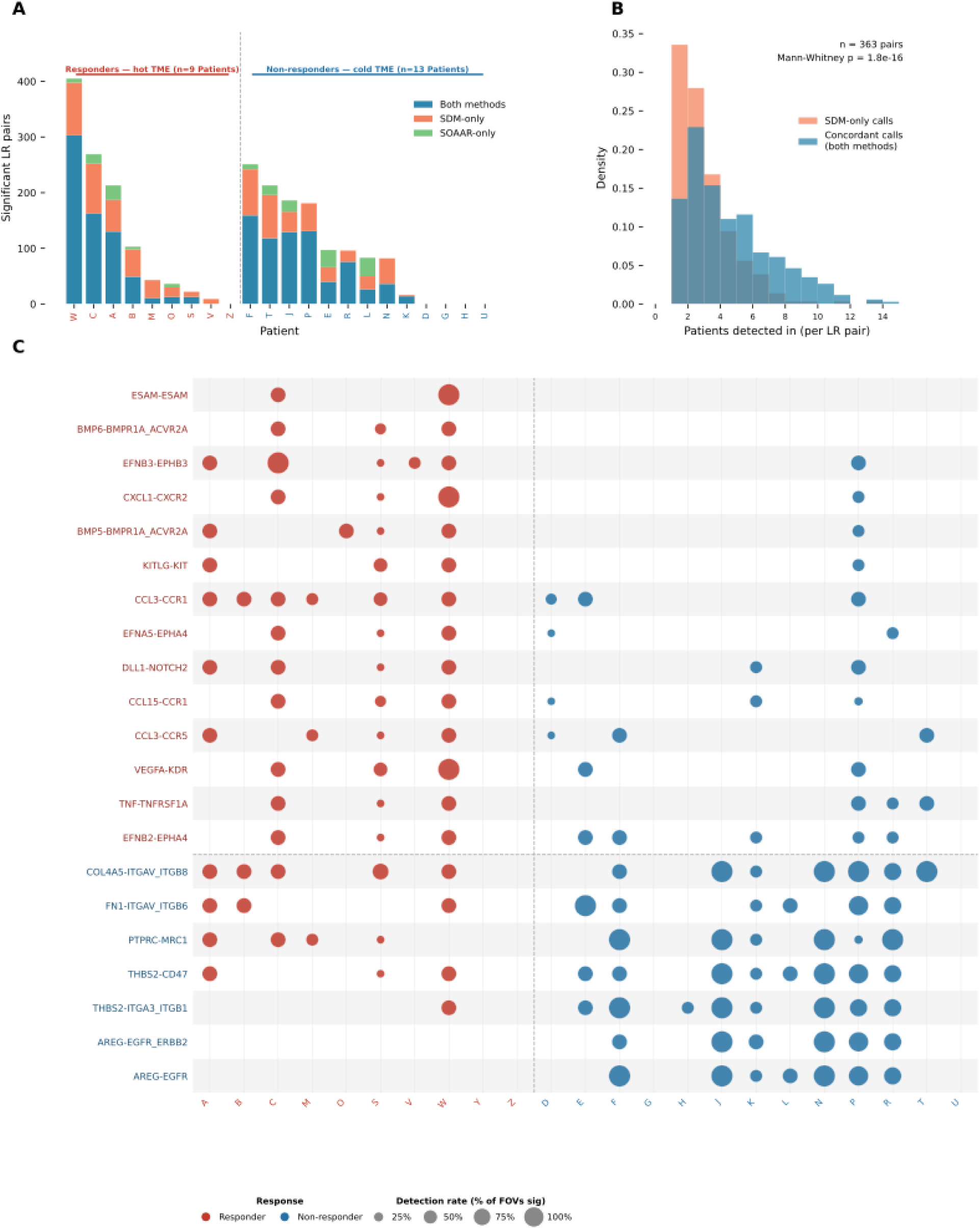
SOAAR identifies immune cell ligand-receptor interactions in single-cell spatial transcriptomics CosMX SMI data from HNSCC patients treated with immunotherapy. A. Total number of significant ligand-receptor (LR) pairs detected per patient sample across 9 responders (red, hot TME) and 13 non-responders (blue, cold TME), stratified by method concordance: interactions detected by both SOAAR and SpatialDM (blue), SpatialDM only (orange), and SOAAR only (green). B. Distribution of the number of patient samples in which each LR pair was detected as significant, stratified by method concordance. Interactions called by both methods (blue) were detected in significantly more patients than those called by SpatialDM alone (orange; Mann-Whitney p = 1.8×10⁻¹□, n = 363 pairs). C. Dot plot of selected LR interactions enriched in responders (red, top) or non-responders (blue, bottom) across all 22 patient samples. Dot size indicates the proportion of fields of view (FOVs) in which the interaction was deemed significant for that patient; dot color indicates response group.

With more samples and higher resolution, we identified immune interactions that shape immunotherapy response and resistance in head and neck cancer (**Fig. 4C, Table S2, S3**). Expression of genes involved in the BMP5–BMPR1A_ACVR2A interaction has been associated with improved overall survival in breast cancer [29]. The interactions CCL3-CCR1, CCL15-CCR1, and CCL3-CCR5 have been observed to facilitate T cell infiltration and improve immunotherapy efficacy [30, 31]. DLL1-NOTCH2 interactions have been seen as driving increased CD8 T cell toxicity and the polarization of macrophages towards a pro-inflammatory state [32]. The ITGAV interactions identified in non-responsive patients are associated with canonical extracellular matrix remodeling as a resistance mechanism, a mechanism documented in other cancer types [33]. The interaction between PTPRC and CD206 has been established as a marker of macrophage-mediated immunosuppression, with prior work identifying CD206 as an immunosuppressive macrophage marker [34, 35]. The interactions involving THBS2 both limit immune infiltration and expand cancer cell invasiveness, thereby establishing an environment that reduces immunotherapy efficacy [36, 37]. Finally, the AREG-EGFR interactions drive sustained cancer cell proliferation, in line with the EREG-EGFR interactions observed in the non-responsive Visium samples [38].

## DISCUSSION

Existing widely used ligand-receptor interaction analysis methods for spatial omics data, such as SpatialDM [12] and Meringue [13], adapt bivariate Moran’s I to identify statistically significant interactions but rely on permutation techniques based on heuristics from single-cell analysis. This failure to account for univariate single structure leads to increased false-positive rates, as shown in our simulation studies. To address this, we have developed SOAAR to account for spatial autocorrelation in the permutation procedure through a Moran Spectral Randomization-based approach. SOAAR is the first adaptation of Moran Spectral Randomization to hypothesis testing in spatial transcriptomics.

SOAAR preserves gene-wise autocorrelation in resampled datasets and, as a result, lowers false-positive rates, as seen in the simulation studies. Both of these properties arise from the preservation of spectral power density across the Moran Eigenvectors. While SOAAR has a lower nominal power, when calibrated, its performance is comparable to SpatialDM’s analytic z-score. While they are equally sensitive under calibrated power, in a real-world setting calibration requires a priori knowledge, which is seldom available, meaning that the more calibrated statistical test of SOAAR is preferable for unknown data, as it will detect fewer spurious correlations. Critically, the marginal distributions of the simulated genes are standard normal, matching the distributional assumption underlying SpatialDM’s analytic calculation and showing that failure to account for spatial autocorrelation increases the false positive rate.

The results of the simulation carry over to our analysis of HNSCC Visium data, where we can clearly see that SOAAR identifies fewer statistically significant interactions across samples. Due to the low number of interactions unique to SOAAR, our analysis focused on comparing interactions concordant between SOAAR and SpatialDM with those that SpatialDM identifies uniquely. The concordant interactions are shared across more samples and generally have higher Moran’s I within each sample, indicating that SOAAR’s more calibrated statistical test can also filter low-effect-size interactions. Our analysis identifies responder- and non-responder-enriched interactions that may serve as potential biomarkers of immunotherapy response. The immune interaction analysis of the HNSCC CosMx data provides additional support for the identified responder- and non-responder-enriched interactions in spatial omics data. It shows that an immune-infiltrated, inflammatory environment associated with effective immunotherapeutic intervention, while the prevalence of immunosuppressive interactions and fibrotic extracellular matrix remodeling limits the efficacy of immunotherapy. Overall, SOAAR’s calibrated analysis enables comprehensive cell-cell interaction analysis for individual complex tissue samples while also identifying broader patterns across samples and platforms.

Although SOAAR identified potential LR interactions associated with effective immunotherapy reported in the literature, validation in a larger, independent cohort is warranted. In addition, we found enrichment of traditionally considered pro-tumoral interactions in the Responders, such as VEGFA-KDR, CXCL1-CXCR2, and KITLG-KIT. However, these interactions may indicate a controlled inflammatory state that drives immune infiltration and provides the foundation for a successful response to immunotherapy. These findings provide hypotheses for future investigation and validation in larger patient cohorts.

SOAAR has limitations that warrant future development. First, spatially resolved transcriptomics data are sparse and discrete prior to continuous transformation and z-scoring, which are requirements for a measure like Moran’s I [39]. However, even after transformation, their distributions are not normally distributed, which limits the statistical power of Pearson-like correlation measures such as Moran’s I [40]. Alternative spatial correlation measures could be used instead of bivariate Moran’s I to account for this while still utilizing the underlying phase randomization framework employed by SOAAR. Second, the global nature of the Moran Eigenvectors means that it defines an orthonormal basis on the entire spatial slide but not within specific spatial neighborhoods, which limits the ability to test local Moran’s I for ligand-receptor since expression level localization is independently moved for each gene. Application of a joint rotation across constituent genes of a ligand-receptor interaction may allow this testing to be done more effectively, along with a test statistic such as a spatial scan statistic in a region containing a given location, rather than a test statistic for that individual location [41]. Linking ligand-receptor interactions to downstream intracellular pathways could help elucidate the role of intercellular signaling in prompting and shaping intracellular activity. Future work to extend local SOAAR with the recent GIS-ROTA method for spatial pathway analysis would provide a starting point for such analysis [42]. The need for spatially aware inference goes beyond ligand-receptor analysis. Emerging methods in spatial biology, such as perturbation-enrichment analysis for spatial CRISPR screens, have identified spatial autocorrelation as a potential limitation of permutation-based statistical tests and calls for methods that account for it explicitly [43]. SOAAR offers one possible approach, and the underlying framework is not specific to testing bivariate Moran’s I – it provides a general way of generating spatially constrained null distributions where permutations-based tests are applied.

## CONCLUSION

Here, we provide a hypothesis-testing framework for spatial correlations in spatial omics data motivated by spatial statistics and spectral graph theory. In simulations, SOAAR can create surrogates that preserve gene-wise spatial autocorrelation while reducing false positives that arise from spatial autocorrelation-driven confounding. In a real data example, SOAAR filters out low-signal interactions, mostly in fewer samples, without eliminating interactions with strong signals present in more samples, providing confidence to prioritize future validation work.

## METHODS

### Statistical testing framework

SOAAR tests whether two spatially resolved genes X and Y are significantly co-localized given their individual spatial autocorrelation structures. To do so, SOAAR draws surrogate datasets X∗ by applying Moran Spectral Randomization (MSR) to X, generating spatial arrangements that preserve power spectral density between X and each eigenvector (singleton procedure) and X and each eigenvector pair (pair procedure). In the case of the singleton procedure and the sequential pair procedure, the univariate spatial autocorrelation of X* approximately matches the observed univariate spatial autocorrelation 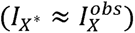. Because Y is held fixed at its observed values, each surrogate produces a bivariate Moran’s I between X∗ and Y, denoted as I_X*Y_.

### SOAAR’s Hypothesis Testing

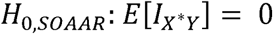

where the expectation is taken over surrogate datasets X* whose univariate spatial autocorrelation approximately matches the observed value 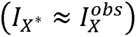, a consequence of minimal spectral redistribution between eigenvectors under the singleton and sequential pair procedures.

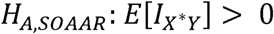

under the same constraint.

### Exchangeability justification

Under H_O,SOAAR_ — spatial independence of X and Y — any spatial arrangement of X with autocorrelation 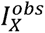 is equally likely to have been the observed configuration. Each surrogate X* is a draw from this constrained null, so the surrogate statistic I_X*Y_ and the observed statistic 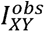 are exchangeable under H_O,SOAAR_. The one-sided p-value is therefore:

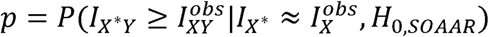

which is estimated as the proportion of nperms surrogates producing a bivariate statistic at least as large as the observed value. This exchangeability holds conditionally on 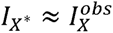 and Y fixed, and does not require X to be spatially random, distinguishing SOAAR from the other methods described below.

Other methods’ (e.g. SpatialDM) Hypothesis Testing

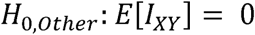

where the expectation is taken over surrogate datasets in which both genes are spatially random (I_X_ ≈ 0 and I_Y_ ≈ 0).

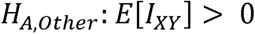

under the same constraint.

### Comparison of Hypothesis Testing Frameworks

For other methods, spatial independence of X and Y implies that any location relabeling of both genes jointly is equally likely, so the joint permutation statistic *I_XY_* is exchangeable with 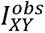 unconditionally. This exchangeability holds only when *I_X_* ≈ 0 and *I_Y_* ≈ 0, making the null calibrated solely under spatial randomness of both genes. When genes are spatially autocorrelated, other methods sample from an exponentially larger region of permutation space than the constrained subset that SOAAR targets, underestimating the null variance and inflating false-positive rates. SOAAR’s conditional exchangeability is valid for any value of 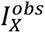, and the two approaches coincide when 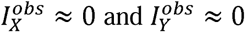.

Due to its calibrated nature, SOAAR also facilitates downstream comparison of multiple samples from similar tissue with different clinical outcomes to identify spatial interactions that separate outcomes, without inflating low-signal interactions. Accounting for the spatial autocorrelation of the tested genes enables adaptive variances for the bivariate Moran’s I of gene pairs based on the autocorrelation regimes of the constituent genes.

### Building spatial weight matrices

Two weight matrices are constructed by default to separately model short-range (cell-ECM and cell-cell contact) and long-range (secreted-signaling) interactions:

- *Short-range weights* (W_short_) are constructed as a radius-nearest-neighbors graph with radius r = d_max_, where d_max_ is the maximum spot-to-spot distance among adjacent spots (computed as the median nearest-neighbor distance × √2 for Visium hexagonal arrays). Entries are binary:

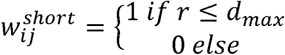

- *Long-range weights* W_long are constructed using a radial basis function kernel: 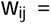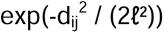 with bandwidth ℓ = 2·d_min_, where d_min_ is the minimum spot-to-spot distance among adjacent spots. The diagonal is set to zero (w_ii = 0) when applied to single-cell resolution spatial omics data.

### Eigendecomposition

For each weight matrix W, SOAAR computes the doubly centered matrix C = HWH, where H = I_n_ − (1/n)11 is the centering matrix, then computes its eigendecomposition C = MΛM with eigenvectors sorted by descending eigenvalue. For datasets with n ≤ 5000, the full eigendecomposition is computed using scipy.linalg.eigh. For larger datasets, a Nyström approximation is used: n_landmarks_ = min(2000, n) landmark points are selected by farthest-point sampling on the spatial coordinates, the eigendecomposition is computed on the landmark subgraph, and approximate eigenvectors are extended to the full sample via the Nyström formula. The n_components_ parameter controls how many eigenvectors are retained for downstream randomization; by default, all are retained for n<=5000, and the top and bottom 1000 are retained for n>5000.

### Spectral randomization

For a given feature vector z (standardized to zero mean and unit variance), spectral coefficients are computed as c = MDz. Surrogates are generated by one of three procedures, selected via the randomization parameter:

- Singleton (randomization="singleton"). Each coefficient is multiplied by an independent random sign: c* = s* · c, with s ∼ Uniform({−1, +1}).
- Sequential pair (randomization="sequential_pair", default). Eigenvectors are grouped into adjacent pairs (2k−1, 2k) for k = 1, …, ⌊n/2⌋. Within each pair p, the amplitude 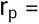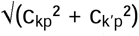 and phase θ_p_ = arctan2(c_k’p_, c_kp_) are computed, then the phase is rotated by D_p_ ∼ Uniform[0, 2π) independently across pairs: 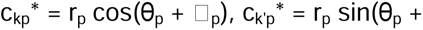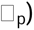. If n is odd, the final eigenvector is treated as a singleton.
- Random pair (randomization="random_pair"). As above, but pairs are formed by random matching of eigenvector indices rather than adjacency. A new random pairing is drawn for each surrogate.

The surrogate feature vector is reconstructed as z* = Mc*, then rank-mapped to the empirical distribution of the original z to restore the marginal: z_final_*[i] = z[argsort(z)[rank(z*)[i]]]. The rank-matching step preserves the original feature’s empirical marginal exactly but introduces small departures from exact preservation of spatial autocorrelation due to the non-linearity of rank matching. In asymmetric mode (mode="asymmetric", default), only the first feature of each pair is randomized while the second is held fixed. In symmetric mode (mode="symmetric"), both features are randomized independently.

### Test statistic and p-value computation

For each feature pair (x, y) with weight matrix W, the observed bivariate Moran’s I is computed as I_obs_ = (n / S_0_) · (z□□ W z□) / √((z□□ H z□)(z□□ H z□)), where S_0_ = Σ□□ w□□. A null distribution is generated from n_perms_ surrogates (default: 999) and a one-sided right-tailed p-value is computed as p = (#{I* ≥ I_obs_} + 1) / (n_perms_ + 1). Multiple-testing adjustment across all tested pairs is performed using the Benjamini-Hochberg procedure with statsmodels.stats.multitest.multipletests. Significance is reported at an adjusted p < 0.05 by default; the alpha parameter defines the significance threshold.

### SOAAR Output

Results are written into adata.uns["soaar_results"] as a DataFrame with columns: feature_1_, feature_2_, interaction_type, I_obs_, pvalue, pvalue_adj, significant. The null distribution per pair is optionally retained (store_null=True) in adata.uns["soaar_null"] for downstream diagnostics. Results may be exported via soaar.export_results(adata, path).

### Simulation Studies

Each of our simulation studies was performed using the following procedure, with different sets of parameters, numbers of simulated genes, and methods employed. For each simulation, we simulated genes with standard-normal marginal distributions on a 20x20 square lattice with rook connectivity, and shortest-path distances between all cells of the lattice were calculated using the Floyd-Warshall algorithm. Using the matrix exponential of the autocorrelation parameter, we then define a pairwise covariance matrix between lattice cells with geometric decay. We then applied the Cholesky decomposition to the pairwise covariance matrix and multiplied it by a vector x ∼ Normal(0, I_400_) to generate a spatially autocorrelated gene expression vector. Since the simulated genes have normally distributed marginals, we do not use the rank-matching step we apply for real data.

To evaluate the ability of SOAAR and random permutations to preserve spatial autocorrelation, we used 39 autocorrelation coefficients ranging from -0.95 to 0.95 in increments of 0.05. We then used SOAAR’s singleton randomization, sequential eigenvector pair randomization, random eigenvector pair randomization, and a uniform random vector randomization approach to evaluate the root mean squared error of the spatial autocorrelation of 1,000 surrogates per simulated gene.

To test SOAAR’s ability to account for spatial autocorrelation in bivariate spatial correlation testing and calling false positives at a significance of 0.05, we composed a grid of pairs of spatial autocorrelation coefficients using the following logic:

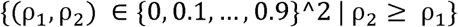

For each pair of coefficients, we generated 10,000 simulated gene pairs with a spatial cross-correlation between the genes of 0. We then used the same data for both SpatialDM’s random permutation procedure and its analytic p-value procedure, which assumes that both genes are normally distributed and that their expression patterns are spatially random. Because SpatialDM requires non-negative input expressions, all simulated datasets were shifted by adding the absolute value of the minimum expression for each gene. This preserves the variance and yields the z-transformed variable used in Moran’s I calculation, matching the original simulated expression used in SOAAR’s calculation. To evaluate SOAAR here, we used the sequential eigenvector pairing procedure.

To test statistical power, here we generated 75 triples of two spatial autocorrelation coefficients and a bivariate cross-correlation coefficient as follows:

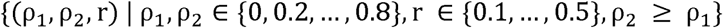

For each triplet, we generated 1,000 pairs of simulated genes. As in the false-positive simulation, we apply SpatialDM’s permutation procedure and its analytic p-value calculation to simulated datasets that are shifted by the minimum expression for each gene. We apply SOAAR’s sequential eigenvector pairing procedure.

### Data used and preprocessing

Visium data of 10 HNSCC patients treated with immune checkpoint blockade therapy were downloaded from NCBI GEO (ID: GSE301720) [16, 44]. Genes in the HNSCC Visium datasets that were expressed in fewer than 3 spots were filtered out. Each spot’s expression vector was normalized by dividing by the total number of reads and then multiplying by the median total reads across all spots. The data was then log1p transformed. The ligand-receptor interaction database CellChatDB [2] was used as the data source for ligand-receptor interactions, multimeric complex identification, and interaction type determination.

To identify immune interactions from CosMX SMI data (https://datadryad.org/dataset/doi:10.5061/dryad.n2z34tn7x), we filtered FOVs to include only those with 50 or more immune cells annotated. Our analysis focused on interactions in which immune cells were either the sender or the receiver. Given that some patients had multiple FOVs, we linked an interaction to a given patient if it was statistically significant in at least half of the FOVs from that patient’s tissue biopsies.

### Software dependencies

SOAAR is implemented in Python 3.10+ and depends on numpy ≥ 1.24, scipy ≥ 1.10, scikit-learn ≥ 1.3, anndata ≥ 0.10, pandas ≥ 2.0, and statsmodels ≥ 0.14.

## Data and Code Availability

SOAAR can be installed from GitHub (https://github.com/pkhatri94/SOAAR) or PyPI. Documentation and tutorials are available on SOAAR.readthedocs.io.

## Author contributions

P.K and H.Q.D conceptualized the study. P.K was responsible for method development and implementation, with input from H.Q.D. and C.K. Analysis was conducted by P.K, with contributions from H.Q.D. The initial draft of the manuscript was written by P.K. and H.Q.D., with review and editing by M.A.N and C.K. Supervision was provided by H.Q.D. and C.K.

---

**Figure S1.**
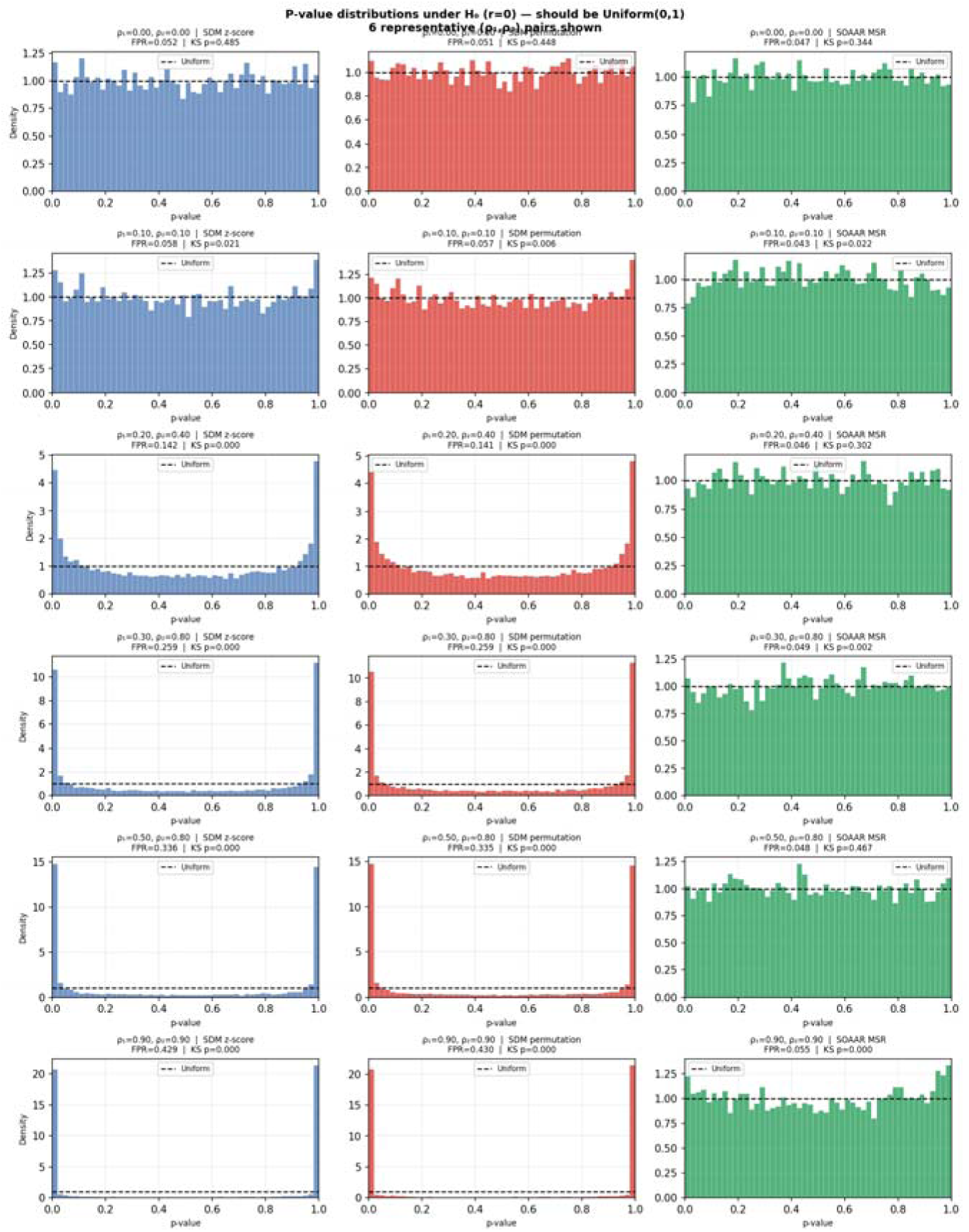
P-value histograms for SpatialDM z-score (blue, left column), SpatialDM permutation (red, center column), and SOAAR MSR (green, right column) across six representative pairs of spatial autocorrelation coefficients (ρ₁, ρ₂), evaluated on 10,000 simulated null gene pairs per coefficient pair (r = 0) on a 20×20 rook grid. A. At no autocorrelation (ρ₁ = ρ₂ = 0), all three methods produce approximately uniform p-value distributions with FPR near 0.05. B. At mild autocorrelation (ρ₁ = ρ₂ = 0.1), SpatialDM’s methods show a slight increase in FPR, while SOAAR remains well-calibrated. C. At moderate autocorrelation (ρ₁ = 0.2, ρ_2_ = 0.4), SpatialDM methods show clear false positive inflation, while SOAAR remains close to uniform. D. At even higher autocorrelation (ρ₁ = 0.3, ρ_2_ = 0.8), SpatialDM methods show severe false positive inflation, while SOAAR shows minor deviation from uniformity. E. At (ρ₁ = 0.5, ρ_2_ = 0.8), SpatialDM inflates false positives even further while SOAAR retains its calibration. F. At the most extreme autocorrelation regime (ρ₁ = ρ₂ = 0.9), SpatialDM’s method reach a false positive rate over 8 times that of the empirical false positive rate while SOAAR shows only a moderate degree of inflation.

Supplementary Table 1 – Combined results across all 10X Visium datasets for all tested ligand-receptor interactions with the following fields:

- Sample: The sample ID
- Response_group: Response status to immunotherapy
- Ligand: Ligand identity
- Receptor: Receptor identity
- Regime: Secreted signaling/contact signaling/ECM interaction label
- Concordant: Called significant by both SpatialDM and SOAAR
- sdm_only: Called significant by only SpatialDM
- soaar_only: Called significant by only SOAAR
- neither: Called significant by neither SpatialDM nor SOAAR
- soaar_sig: Called significant by SOAAR (soaar_fdr < 0.05)
- soaar_pval: uncorrected p-value based on SOAAR’s test
- soaar_fdr: BH corrected p-value based on SOAAR’s test
- z_score: z-score of SOAAR p-value based on null mean and variance calculated by SOAAR
- sdm_sig: Called significant by SpatialDM (sdm_fdr < 0.05)
- sdm_pval: uncorrected p-value based on SpatialDM’s test
- sdm_fdr: BH corrected p-value based on SpatialDM’s test
- observed: Observed bivariate Moran’s I
- null_mean: Mean bivariate Moran’s I of null distribution
- null_var_msr: Bivariate Moran’s I variance based on SOAAR’s MSR framework
- var_random_null: Bivariate Moran’s I variance based on SpatialDM
- log10_var_ratio: log10 of the variances calculated by both methods
- pearson_lr: Pearson correlation between ligand and receptor

Supplementary Table 2 – Combined results across all CosMx Fields of View for all tested ligand-receptor interactions with immune cells as either senders or receivers with the following fields:

- Ligand: Ligand identity
- Receptor: Receptor identity
- Annotation: Secreted signaling/contact signaling/ECM interactions
- weight_used: Weight matrix used, W_secreted or W_contact
- I_obs: Observed bivariate Moran’s I
- p_soaar: uncorrected p-value based on SOAAR’s statistical test
- p_sdm_z: uncorrected p-value based on SpatialDM’s analytic z-score calculation
- p_sdm_perm: uncorrected p-value based on SpatialDM’s permutation
- immune_mode: Whether an immune cell was the sender or receiver
- fdr_soaar: BH corrected p-value based on SOAAR’s statistical test
- fdr_sdm_z: BH corrected p-value based on SpatialDM’s analytic z-score calculation
- fdr_sdm_perm: BH corrected p-value based on SpatialDM’s permutation
- sig_soaar: Called significant by SOAAR’s statistical test (fdr_soaar < 0.05)
- sig_sdm_z: Called significant by SpatialDM’s analytic z-score (fdr_sdm_z < 0.05)
- sig_sdm_perm: Called significant by SpatialDM’s permutation (fdr_sdm_perm < 0.05)
- run: Tumor microarray name and index
- fov: Field of view index

Supplementary Table 3 – Linking TMA+fov identities from CosMx assay to deidentified patients and their immunotherapy response status

- FOV_ID: run + fov from table 2 concatenated and underscore delimited
- Patient_ID: Deidentified patient, indicated by letter
- Response: Immmunotherapy response status

